# Using Beads as a Focus Fiduciary to Aid Software-based Autofocus Accuracy in Microscopy

**DOI:** 10.1101/2025.03.18.643916

**Authors:** I Gibson, E.J Osterlund, R Truant

## Abstract

Brightfield microscopy is an ideal application for studying live cell systems in a minimally invasive manner. This is advantageous in long-term experiments to study dynamic cellular processes such as stress response. Depending on the sample type and preparation, the inherent qualities of brightfield microscopy, being very low contrast, can contribute to technical issues such as focal drift, sequencing lags, and complete failure of software autofocus systems. Here, we describe the use of microbeads as a focus aid for long-term live cell imaging to address these autofocus issues. This protocol is inexpensive to implement, without extensive additional sample preparation and can be used to capture focused images of transparent cells in a label-free manner. To validate this protocol, a widefield inverted microscope was used with software-based autofocus to image overnight in time-lapse format, demonstrating the use of the beads to prevent focal drift in long-term experiments. This improves autofocus accuracy on relatively inexpensive microscopes without using hardware-based focus aids. To validate this protocol, the KNIME logistics software was used to train a random forest model to perform binary image classification.

**Key features:**

- label-free live cell imaging in time-lapse format
- Troubleshooting software autofocus for brightfield mode

**Graphical overview:** 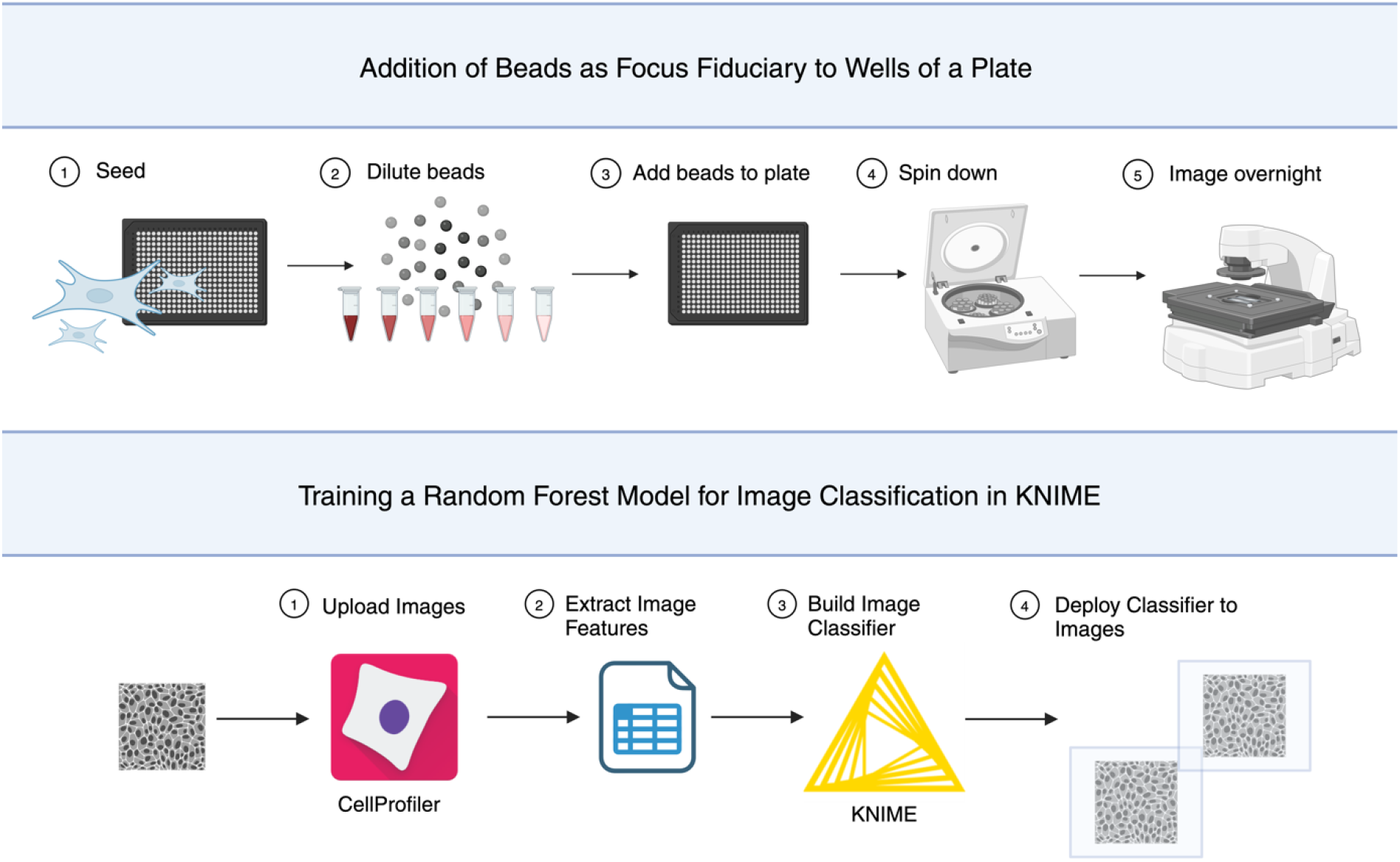

## Summary

In fluorescence microscopy, fluorescent stains and probes can be used to highlight subcellular structures. The caveats of using fluorescent microscopy, such as photobleaching and phototoxicity are well described^1^. Phototoxicity refers to the process where typical wavelengths that are used in fluorescence imaging generate excessive reactive oxygen species (ROS) which can lead to DNA damage^2^. This phenomenon is dependent on wavelength and length of exposure to cells^2^. Intracellular ROS production can be buffered by cell antioxidant molecules and enzymes, although in fluorescent imaging, ROS buffering capacities can be exceeded^3,4^. Photobleaching refers to the destruction of a fluorophore such as a fluorescent protein (i.e., GFP) in the excited state that leads to depletion of signal over time which can result in the production of ROS and subsequent photobleaching^1,5^. Interaction of a fluorophore in the excited triplet state and molecular oxygen may result in the production of reactive oxygen species and cause photobleaching^5^.

Commonly used stains such as DRAQ5 and Hoechst that highlight chromatin have demonstrated cytotoxic effects in live-cell imaging^6,7^. DRAQ5 dye alters the dynamics and localization of critical proteins involved in DNA transcription, replication and repair and Hoechst can cause cell cycle arrest or delay of the G2 phase^6,7^. Consideration must be taken when designing experiments to minimize confounding effects of potentially cytotoxic stains, phototoxicity and photobleaching, especially in long-term acquisitions. This is necessary as artifacts may appear from sample preparation and/or in imaging and this becomes increasingly important when evaluating the effects of treatments to cells.

Time-lapse experiments are easily susceptible to phototoxicity. Cell migration is reduced at high fluorescent light doses compared with cells imaged in brightfield^8^. Time resolution, and/or light dose must be compromised to reduce phototoxicity using fluorescent microscopy, making this unsuitable to follow rapid cellular activities^9^.

In contrast, Brightfield microscopy is label-free, making this technique less invasive and relatively inexpensive. This is an ideal application for studying live-cell systems and processes which are sensitive to oxidative stress caused by elevated ROS such as mitosis^4,10^. Additionally, information which may differ from fluorescent images such as cell texture can be acquired simultaneously with structural information to create novel morphological profiles^11^.

Autofocus combines search algorithms and focus metrics to identify the sharpest image across the entire focal depth^12,13^. In automated microscopy, focus metrics are contrast-based algorithms applied to each image when scanning the focal depth. Plotting the magnitude of the focus function creates a focal curve where the most focused image is the global maximum^12^. Ideally, the range of the focus curve is narrow creating a single defined global maximum with few if any local maximum^10^. In practice, when evaluated across different sample types, commonly used focus metrics are far from robust and the focal curve is not unimodal^14,15,16^.

Confounding the susceptibility of focus metrics to failure, search algorithms are another possible source of error in autofocus. These algorithms represent optimization problems, where the most focused image must be defined accurately and rapidly. Global search is a commonly used search algorithm, which samples the entire focal depth but is computationally slow. To optimize speed, the hill climbing method is a type of binary search using a combination of fine and rough focusing but can incorrectly select a local maximum as the most focused image^16^. The susceptibility to failure of autofocus systems appears frequently in the presence of plate scratches, debris, noise, lack of high-frequency content (i.e., few cells, transparent cells) or uneven illumination^14,15^. High-frequency content is associated with sharp objects. Contrast-based autofocus systems identify the global maximum by comparing sharpness across the focal depth^14^.

Autofocus algorithms have been evaluated and implemented effectively with fluorescence microscopy, due to the inherent high signal-to-noise ratio^17^. Despite mitigating many drawbacks associated with fluorescence microscopy, the qualities of Brightfield microscopy make it more susceptible to autofocus problems. For example, imaging transparent cells such as fibroblasts, produces images with low-signal-to-noise ratio and makes identifying the correct focal plane by autofocus more error-prone.

Our goal was to utilize an inverted widefield microscope with software autofocus to capture mitotic events in hTERT immortalized patient-derived fibroblasts, also known as “TruHD” cells^18^, to study the impact of Huntington’s disease-associated mutations on cell growth. Since increased cell stress and sensitivity to DNA damage have been demonstrated in TruHD cells^18^, we wished to mitigate additional cell stress by choosing to collect brightfield stain-free images. However, we found the software-based autofocus insufficient for use in the transmitted light channel specifically (Figure 1). This failure could not be resolved by imaging in phase contrast, or by selecting an alternative built-in autofocus algorithms. In collecting temporal data over time without the aid of autofocus, we observed drift from the manually set focus over time even while using stains, demonstrating that a working autofocus is necessary to acquire quality movies. In this paper, we describe, in detail, the use of microbeads as an aid for focusing on the transmitted light channel to acquire images of live cells in time-lapse.

**Figure 1:**
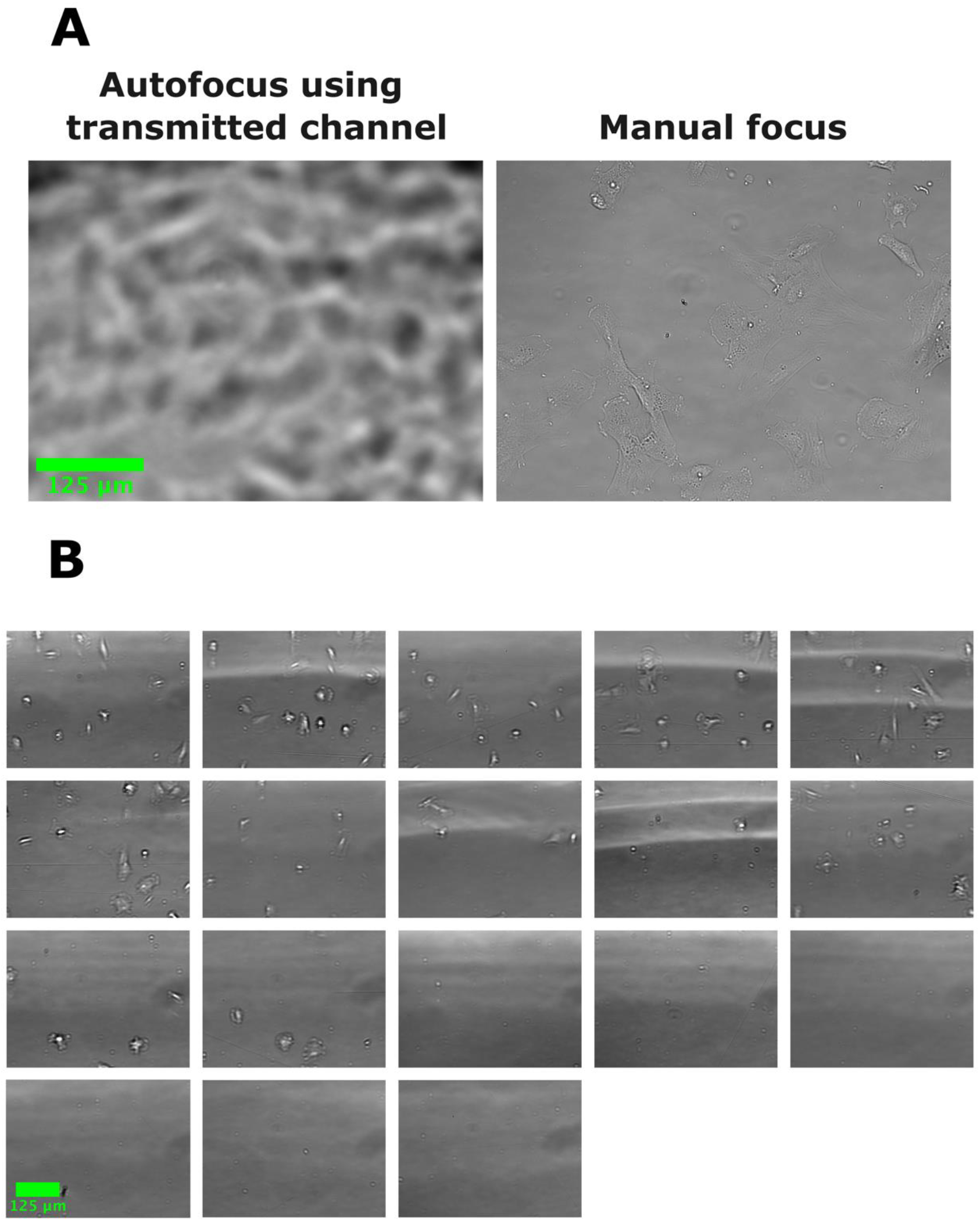
Software Autofocus can fail when acquiring brightfield images. In all images, human fibroblasts were seeded in a 384-well plate and images were acquired with a 20x objective Plan fluorite (Evos, NA=0.5). (A) Example brightfield image acquired using autofocus compared with an image focused manually. Autofocus fails to find the correct plane of focus based on the transmitted light channel (B) Images acquired of other wells of the same plate from (A) in a single automated scan using autofocus. All images show failure of autofocus using the transmitted channel to focus.

This method is not only effective to eliminate long term focus drift, but in multi-well plates, even placing beads in wells adjacent to wells without beads aided focus accuracy. In the place of tedious manual annotation of over 3,700 images collected using this protocol, a random forest model was trained. The model was used to perform unbiased binary classification, identifying images as focused or unfocused. The classifier is not required to use beads as a focal fiduciary, however, this was essential to validate this method. This protocol will briefly describe training the machine learning algorithm using the data analysis platform, KoNstanz Information MinEr, or KNIME (https://www.knime.com/).

## Materials and reagents

### Biological materials

1. TruHD Cells^16^

### Reagents

1. Minimal Essential Media (MEM, 1X) (Gibco, 10370-021)
2. Fetal bovine serum (Wisent, 098450)
3. GlutaMax (Gibco, 35050-061)

### Solutions

1. Supplemented Media (see recipes)
2. Deionized Microfiltered Water

### Recipes

#### 1. Supplemented Media

| Reagent | Final concentration | Quantity or Volume |
| --- | --- | --- |
| Minimum Essential Media 1X | n/a | 500ml |
| Fetal Bovine Serum | 13% | 75ml |
| Total | n/a | 575ml |

### Laboratory supplies

1. 0.2µm filter (Filtropur BT50, 83.3941.101)
2. 50ml Screw cap tube (Starstedt, 62.547.254)
3. Microbeads: **\*\*Beads are stored at room temperature and covered in foil\*\***
  a. Silicon dioxide microparticles 2um (Sigma-Aldrich, Cat No. 81108-5ML-F)
  b. Red fluorescent latex beads 2um (Sigma-Aldrich, Cat No. L3030-1ML)
  c. Non-fluorescent latex beads 3um (Sigma Aldrich, Cat No. LB30-2ML)
4. 384 well plate (PerkinElmer, PhenoPlate)
5. Hemocytometer (any)
6. “Dummy” plate (any)
7. Pipettes tips
  a. 10 µl (VWR, Cat No. 76322-528)
  b. 200 ul (VWR, Cat No. 76322-150)
  c. 1000 ul (VWR, Cat No. 76322-154)
8. Pipettes:
  a. p10 (Eppendorf, Cat No. 3123000020)
  b. P200 (Eppendorf, Cat No. 3123000055)
  c. p1000 (Eppendorf, Cat No. 3124000121
9. 1.5ml Microcentrifuge tubes (Avantar, Cat No. 20170-038)
10. Tissue Culture Plates (Fisherbrand, Cat No. FB012924)
11. Microtube rack (Any)

### Equipment

1. Multi-gas Incubator (PHCBI, MCO-170M-PA)
2. Evos M7000 Microscope (ThermoFisher, Cat No. AMF7000)
  a. EVOS Onstage Incubator (Invitrogen, Cat No. AMC1000)
  b. EVOS Objective lens (20x Air, Plan fluorite 20x/NA 0.5, Cat No. AMEP4698)
  c. EVOS Light Cube (Cy5, Cat No. AMEP4956)
3. Laboratory plate centrifuge (Eppendorf, Centrifuge 5810R)
4. Biological safety cabinet (Microzone corporation, BK-2-4)

### Software and datasets

1. Python (3.10.9)
2. EVOS imaging software (2.1.677.717)
3. GraphPad Prism (10.2.3)
4. KNIME (5.2.1)
  - System requirements:
    i. Linux: 64bit, at least 8gb RAM
    ii. Mac: 10.11 and above and at least 8gb RAM
    iii. Windows: 32 or 64bit and at least 8gb RAM

### Procedure

#### A. Day 1: Plate Preparation

**\*\*see General note 1\*\***

**Critical:** this protocol describes experiment 2 (figure 2), 2um red latex beads are used.

**Figure 2:**
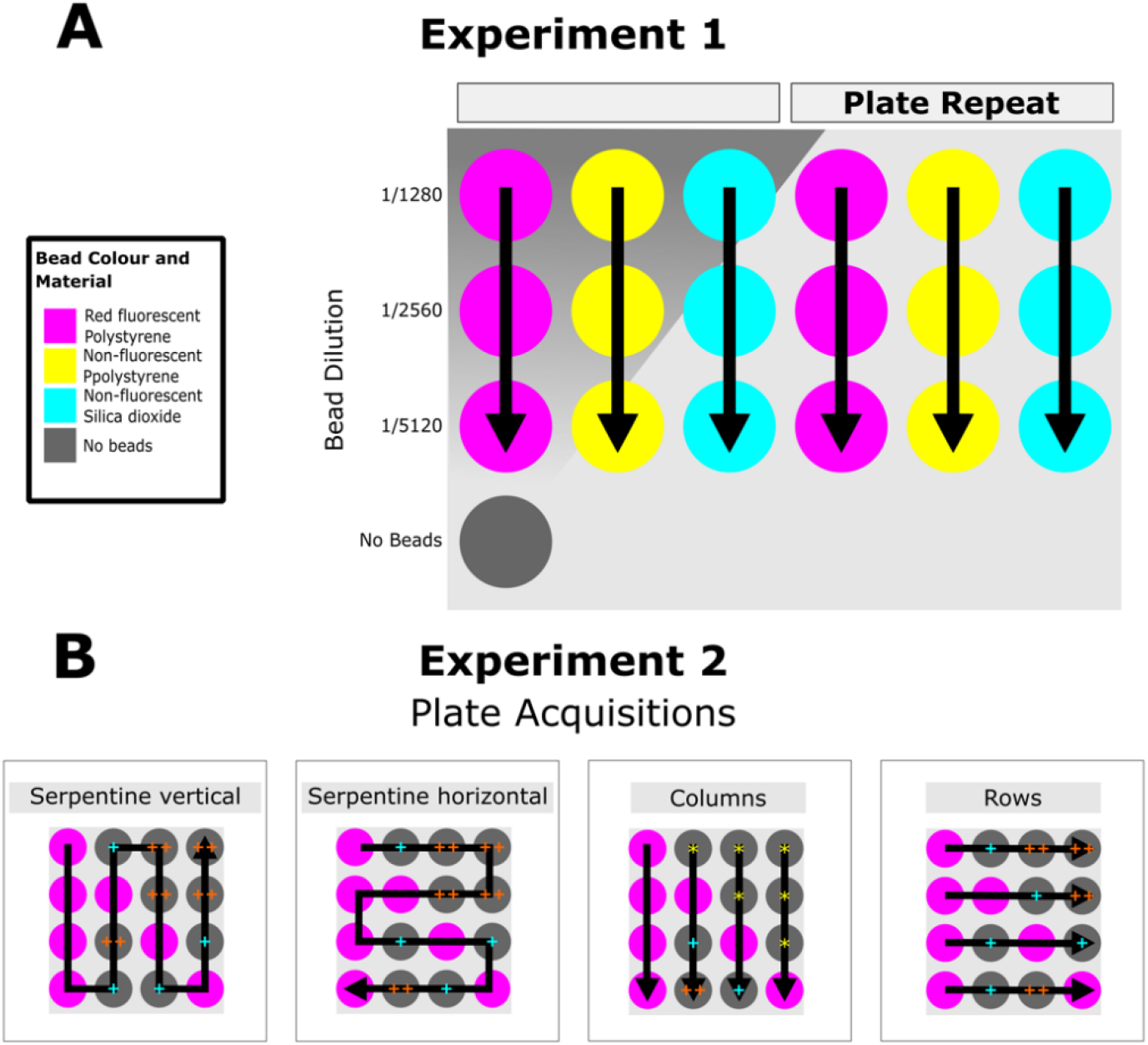
Graphical Schematic of the experimental protocol used for validation. Images analyzed later in this text were acquired using 2 different experimental protocols set up on an EVOS M7000. In both A and B, black arrows indicate the directionality of the selected scan protocol and the color of the well indicates well content (see legend in A). For further details of the autofocus procedure used, see the data acquisition section. **(A)** Experiment 1 was used to evaluate the efficacy of different bead material and fluorescence using polystyrene (PS) 2μm, PS 3μm, Silica Dioxide (glass) 2μm. **(B)** Experiment 2 was designed to determine whether the presence of beads in the well, or in a neighboring well, reduces imaging lags and focus drift. Red fluorescent beads (2um) were used. The transmitted channel was selected as the focus channel and the single channel acquired. The same sample was imaged in quadruplicate using 4 different scanning routings: Serpentine vertical, Serpentine Horizontal, Column wise and Row wise. Wells marked with a yellow asterisk highlight wells where images were acquired before a well containing microbeads; A blue plus sign highlights wells that were imaged directly following (adjacent) a well containing microbeads; 2 orange plus signs highlight wells that were imaged within 2 or more wells following a well containing microbeads. Plate metadata was extracted from all four plate scans, providing the time each image was acquired. The exposure time per well was determined as the time between the first and last frame taken in each well. Subsequently, data was sorted based on the presence or absence of beads in the well images. Wells that did not contain beads were further sorted into 3 categories based on the order imaged (before or after imaging beads) and their relative position in the plate (1 or 2+ wells away from a well with beads).

1. Cell Counting
  a. Add 300ul of suspended cells to a screw cap tube
  b. Add 3ml of supplemented media to the tube
  c. Mix thoroughly
  d. Add a small amount of the mixture (5ul) to a hemocytometer
  e. Follow directions provided for hemocytometer to determine cell dilution
  f. Dilute the cells to 15cells/ul using supplemented media
2. Seed 384 well plate
  a. Add 50ul of the diluted cells from step 1 to each well as shown in experiment 2 (figure 2)
  b. Leave the plate at room temperature for 15 minutes to allow cells to settle on the bottom of the well
  c. Place the plate into the incubator overnight (37.0°C, 5.0% CO_2_, 8.0% O_2_)

#### B. Day 2: Bead Dilution in Filtered Water

**Critical:** cells should be 40-50% confluent

**\*\*see General note 2 and 3\*\***

1. Filter deionized water using a 0.2 µm sized filter
2. Label Microcentrifuge tubes
  a. Place 6 microcentrifuge tubes into the microtube rack
  b. Label with dilutions as follows: (1/11, 1/352, 1/704, 1/1408)
3. Add filtered water to tubes as follows:
  a. Add 50 µl water to 1/11
  b. Add 155 µl water to 1/352
  c. Add 100 µl water to all remaining tubes (from step 2)
4. Serially dilute beads
  a. Add 5 µl beads to 1/11 tube
  b. Mix thoroughly by pipetting at least 10 times
  c. Add 5ul of 1/11 to 1/352 tube
  d. Mix thoroughly by pipetting at least 10 times
  e. Add 100 µl of 1/352 dilution to 1/704 tube
  f. Mix thoroughly by pipetting at least 10 times
  g. Add 100 µl of 1/704 dilution to 1/1408 tube

#### C. Day 2: Addition of beads to pre-seeded 384 well plate

1. Dilute the beads in Media
  a. Label a new microcentrifuge tube as 1/1408M
  b. Add 200ul of supplemented media to the 1/1408M tube
  c. Add 25ul of the beads from 1/1408 dilution to 1/1408M
  d. Aliquot 275ul of supplemented media to a new microcentrifuge tube (25 µl per well is the minimum volume)
2. Add the beads to the plate **\*\*See General Note 4\*\***
  a. From 1/1408M, add 25 µl of the beads to bead wells as shown in experiment 2 (figure 2) to achieve a final dilution of 1/38,016.
  b. Add 25 µl of aliquoted media to wells without beads as shown in experiment 2 (figure 2)
3. Spin down the plate
  a. Place the plate in the plate centrifuge for 5 minutes at 1000 x G
  b. **Critical:** Carefully remove the plate from the centrifuge, avoiding tilting and sudden movements **\*\*Disrupting the beads after spinning the plate will cause them to re-suspend in the media. Avoid any sudden movements when transferring the plate to the microscope and handling after this step\*\***

#### D. Day 2: Collecting Image Data

4. Start the EVOS M7000 system
5. Insert a “dummy” plate onto stage to fill the space while the microscope is warming up
6. Fill the incubator system with water
7. Open the CO_2_ tank for delivery to the sample
8. In the EVOS software, manually activate the environmental controls (5% CO_2_, 37°C)
9. Allow the system to warm up for 1 hour before use
10. Transfer the plate to the microscope
  a. Data Acquisition:
    i. Experiment 1 (Figure 2): **\*\*see General note 5 and 6\*\***
      1. Imaging frequency and protocol: Serpentine vertical scanning, single passage of the plate and 4 fields per well.
      2. Autofocus settings: Algorithm selected: fluorescence optimized, autofocus with all channels, every field, first time point only, skip fields that fail
      3. Objective/channels acquired: 20x Plan fluorite, Transmitted and Cy5
    ii. Overnight Scanning Procedure (Experiment 2, Figure 2):
      1. Imaging frequency and protocol: Serpentine vertical scanning, acquire plate every 4 minutes for 14 hours and 4 fields per well.
      2. Autofocus setting: Algorithm selected: fluorescence optimized, autofocus every field using transmitted channel, every time point, without skipping fields that fail.
      3. Objective/channel acquired: 20x Plan fluorite, Transmitted

## Data analysis

Validation of this protocol was carried out with three technical replicates and two biological replicates. The training of the random forest model is briefly described below to validate this protocol. The model is unnecessary to implement beads as a focal fiduciary.

### Image Metadata analysis

Two scripts were created to extract metadata from the images acquired and used to create a CSV file containing the exposure time in each well. Next, this was sorted based on the position of the well. All Scripts used for data analysis can be found on the Truant Lab GitHub at: https://github.com/TruantLab/Py-files-Bio-Protocols-

### Sample time-lapse video creation

Images were selected from experiment 2 (Figure 2) of a well with beads and an adjacent well (1^st^ adjacent). The images were scaled using the mean pixel intensity of each image set, with a Python script also found on the Truant Lab GitHub.

### Image Classification in KNIME

The workflow described in this protocol is available at: https://hub.knime.com/s/fbbMd4SW0FRbecm4.

### Training a Random Forest Model in KNIME

#### Data preparation and feature extraction in CellProfiler

374 brightfield images were selected to generate a training dataset. Images were manually annotated with the ground truth class (focused, not focused). In CellProfiler, each image was filtered using either a Gaussian filter and/or an edge detection filter. From each image, texture features were extracted from the raw image or filtered images and exported to a database file. Features were graphed to identify candidates to train the image classifier in KNIME. Identified features demonstrated some ability to distinguish image class (i.e., focused, not focused) annotated earlier. The features selected include; angular second moment (extracted from raw images), contrast and sum average (extracted from Gaussian-filtered images) and contrast (extracted from images with Gaussian and edge-detection filters applied to images). Features were extracted with a filter size of 170 pixels. A test set, and all deployment data (i.e., images generated from repeats of this protocol) were analyzed through the CellProfiler pipeline and features were extracted to separate database files.

#### Design of a random forest model training architecture in KNIME

The KNIME workflow consists of training data pre-processing and annotation, training, validation, hyperparameter optimization and deployment (Figure 3).

**Figure 3:**
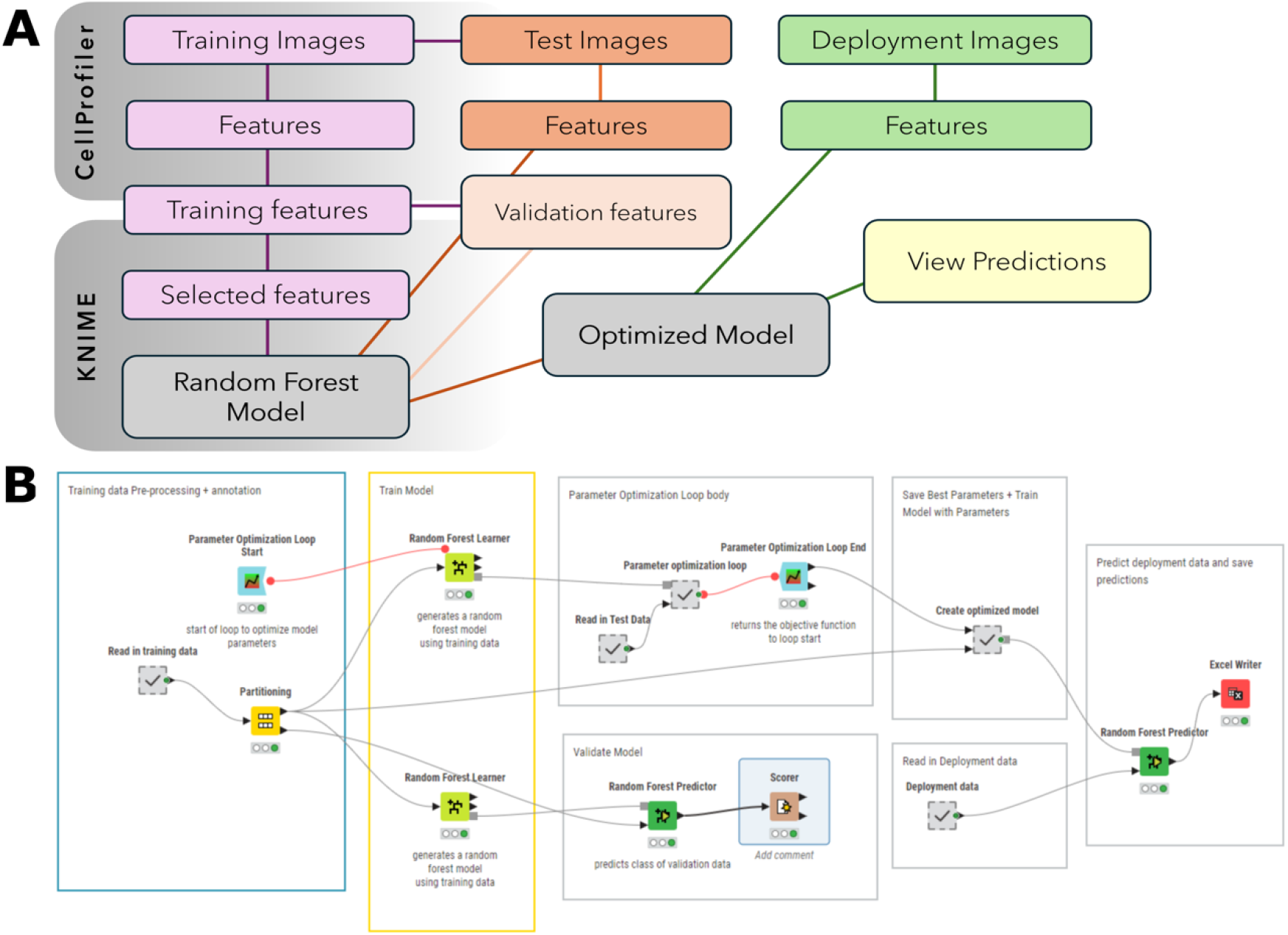
Overview of training a random forest image classifier in KNIME. (A): In CellProfiler image features are extracted from the training, test and deployment images. The features selected for training are read into KNIME and used to train a random forest model to classify images as focused or unfocused. Test features are used to optimize the model. Image features from the deployment images are read into the workflow and classified using the optimized model. (B): Image of KNIME workflow used to train a random forest model. The workflow consists of pre-processing, validation, hyperparameter optimization and deployment. Pre-annotated training data is read into the workflow. Features selected for training are extracted and a second annotation of this data is performed to generate a class variable in KNIME. Training features are divided between validation and training. A loop is used for hyperparameter optimization to determine an appropriate number of models and tree depth to use. The same features selected for training are extracted from the test data and used along with the training data for model optimization. The test set is used to generate the objective value which indicates the performance of each set of hyperparameters as they are evaluated. The best-performing hyperparameters are saved to a table and used to train the deployment model. As before, the same features selected for training are extracted from the deployment data. The classifier is applied to these features and predictions are exported to a spreadsheet file (MS Excel).

#### Probability of a focused Image

Images acquired in experiment 2 (Figure 2) were predicted using the KNIME classifier. The probability of a focused image was calculated as the ratio of focused images to total images acquired in each field.

#### Model validation, optimization and test data

The model performed with 98.23% accuracy with a validation set of the training data. The model had 75% accuracy with the test set before optimization. Following optimization, the model performance improved to 92.97% accuracy with the test set.

## Validation of protocol

To evaluate the use of this protocol two criteria were selected: the presence of imaging lags and the ability to retain focus in overnight image acquisitions as shown by experiment 2 in Figure 2. The utility of microbeads other than that used for the overnight experiments was evaluated as shown in experiment 1 (Figure 2). The effect of adding beads to a well was determined by compiling images in time-lapse format. The image classifier was used to evaluate the effect of the beads to adjacent wells.

### Assessing the consistency in exposure time per well in the presence and absence of beads

Experiment 2 was designed to determine whether the presence of beads in the well or a neighboring well reduces imaging lags. In preliminary experiments, we noticed imaging lags within the first passage of the plate whereby the autofocus feature would expose certain wells longer than others.

The addition of the microbeads promoted consistent exposure time for wells with beads and significantly decreased mean acquisition time as compared to wells before beads (Figure 4). This indicates that the presence of the beads reduced and maintained consistent exposure time across multiple wells. We did not observe a difference to exposure time in wells adjacent to the beads compared to wells before beads. This demonstrates that there is no improvement to the exposure time in adjacent wells. Therefore, having beads in the wells imaged is necessary for consistently low exposure time. However, changes to exposure time per well are not necessarily predictive of image focus. To access this, we determined a means of classifying image focus to further validate the use of this protocol.

**Figure 4.**
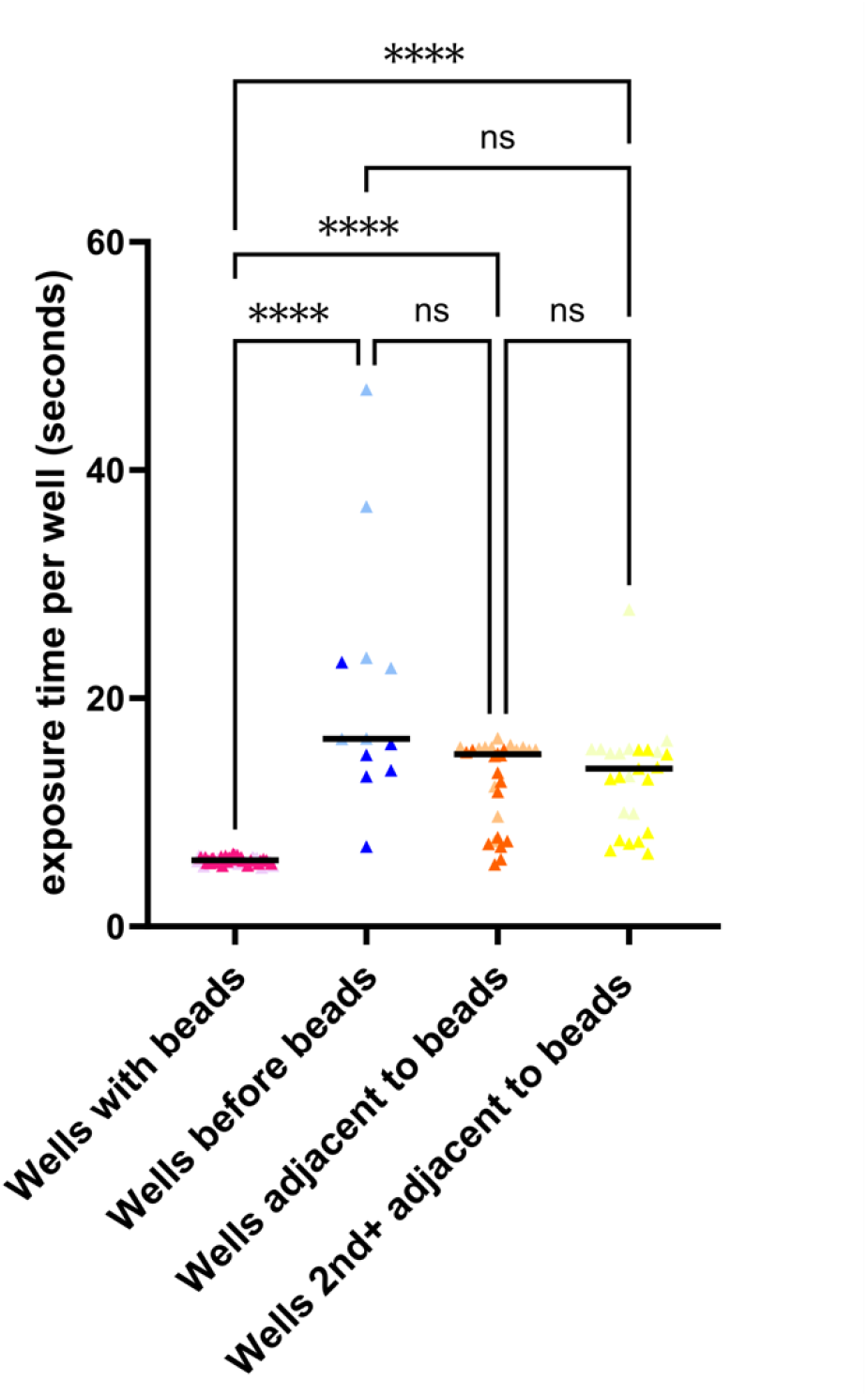
The presence of microbeads in a well reduces exposure time and does not extend to adjacent wells. Images were acquired as described by experiment 2 (see Figure 2B). All analysis excluding graphing was completed in Python. Metadata was extracted from all four plate acquisitions as illustrated in Figure 2B, and the exposure time per well was calculated as the time point difference between the first and last images acquired in each well. Data was separated into; Wells with beads, wells before beads, wells adjacent to beads and wells 2nd+ adjacent to beads. Images acquired of the same well were included in the same or different data sets (i.e., wells with beads) if these came from different plate scans. Kruskal-Wallis Test was performed in GraphPad Prism (10.2.3) (P < 0.0001, ns > 0.05) N = 2.

### Long-term acquisition image quality

Following four plate acquisitions, the plate was imaged in Serpentine Vertical for 14 hours (Figure 2). In the time-lapse movies created from the images, the presence of the microbeads prevented focal drift in the wells with beads (Video 2A). Focal drift was also prevented in a well without beads, directly adjacent to the bead well (Video 2B). To determine if the effect of the beads on adjacent wells was consistent across the entire dataset of images acquired, a machine learning algorithm was trained to classify images as focused or unfocused. The algorithm was trained using a random forest model and the data analysis software; KNIME.

Subsequently, images acquired overnight in experiment 2 were classified using the model. The presence of beads improved the probability of obtaining a focused image compared to images acquired without an adjacent bead well at the first time point (Figure 5A). Initially, it was expected that the presence of the beads would be most effective in retaining focus in directly adjacent wells (i.e., 1st adjacent) and the effectiveness would decrease with distance from the bead well (i.e., 2nd and 3rd adjacent wells). All adjacent wells had a high probability of being focused (Figure 5B). Finally, the presence of beads in adjacent wells was effective in maintaining a high probability of obtaining a focused image in overnight scans (Figure 5C).

**Figure 5:**
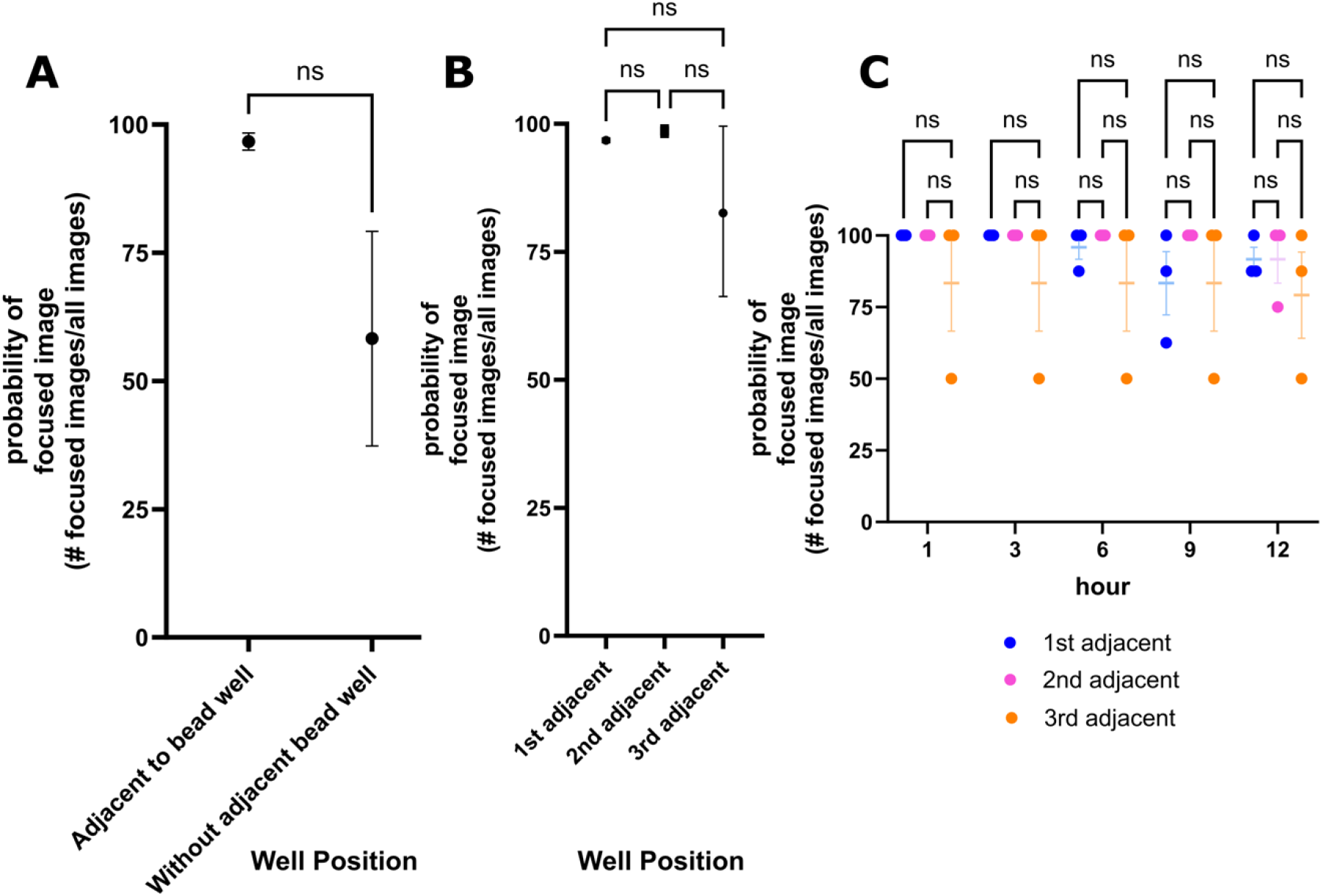
The presence of beads improves the reliability of obtaining a focused image with autofocus. The focus of images acquired in experiment 2 were predicted using the image classifier in KNIME. For a detailed overview of the experimental setup see Figure 2. Each point represents the probability of obtaining a focused image. The likelihood of obtaining a focused image was calculated from the number of focused images acquired and the total number of images acquired and applied to all graphs. (A): The probability of a focused image when beads are scanned before adjacent wells (1st adjacent, 2nd adjacent, 3rd adjacent wells) and when images are acquired without a bead well (n=1). Wells without beads were acquired before wells with beads in experiment 2. 20 Images were analyzed from each repeat, generating a single data point each. The presence of a bead well improves the probability of obtaining a focused image at the first passage of the plate. (B): The position of the well with respect to a bead well and the probability of obtaining a focused image. Images from the complete 14-hour scan were compiled to calculate the probability (n=1). 4440, 4824, and 3216 images are the combined total images acquired across repeats. Image predictions were separated by well position and used to generate a single data point from each repeat. Regardless of the position of the well, obtaining a focused image is highly probable. (C): Hourly assessment of the probability of obtaining a focused image in all adjacent wells (n=1). Images analyzed in panel B were separated by the time point. 48 images were analyzed for each category (i.e., 1^st^ adjacent, 2^nd^ adjacent, 3^rd^ adjacent), representing 8 images analyzed at each timepoint. The presence of beads improves the consistency of acquiring a focused image in a long-term scan. SEM bars are shown, N = 3.

### Efficacy of using distinct types of microbeads as a focus fiduciary

To determine if other beads could be used as a focus aid, fluorescent, non-fluorescent and glass (silicon dioxide) microbeads were tested (Experiment 1, Figure 2A). All channels were used for autofocussing. The non-fluorescent beads appeared more prominently than the cells and exhibited strong autofluorescence (Figure 6). The non-fluorescent beads are larger than both the glass and fluorescent beads and thus, sit further above the plane of the cells as compared to the smaller beads. The objective of adding the beads is to retain focus in long-term scans by preventing focal drift. We hypothesize that if the beads are much larger than the cells, this effect is lost as the autofocus is selecting a plane that is not the optimal focus position. Further, downstream segmentation of these images could be difficult, and the beads may be incorrectly identified as cell objects. The use of silicon dioxide beads produced hazy and low-contrast images compared to the other beads tested. In addition, a caveat of glass beads is that they can focus light by acting as ball lenses. Beads were used at three different dilutions (Figure 6). The effect of the beads to address prolonged well exposure, focal drift and prevent autofocus failures was evaluated at the highest dilution. In preliminary experiments, we found that this allowed for sufficient beads in each field to obtain the intent effect without compromising image appearance. It may be necessary to optimize the bead dilution depending on cell dilution and objective magnification. Evaluation of the effects of the beads on cellular health was limited in validation of this protocol. We did not observe, however, any evidence of interaction between the beads and cells (i.e., phagocytosis) or evident signs of detriment to cellular health.

**Figure 6:**
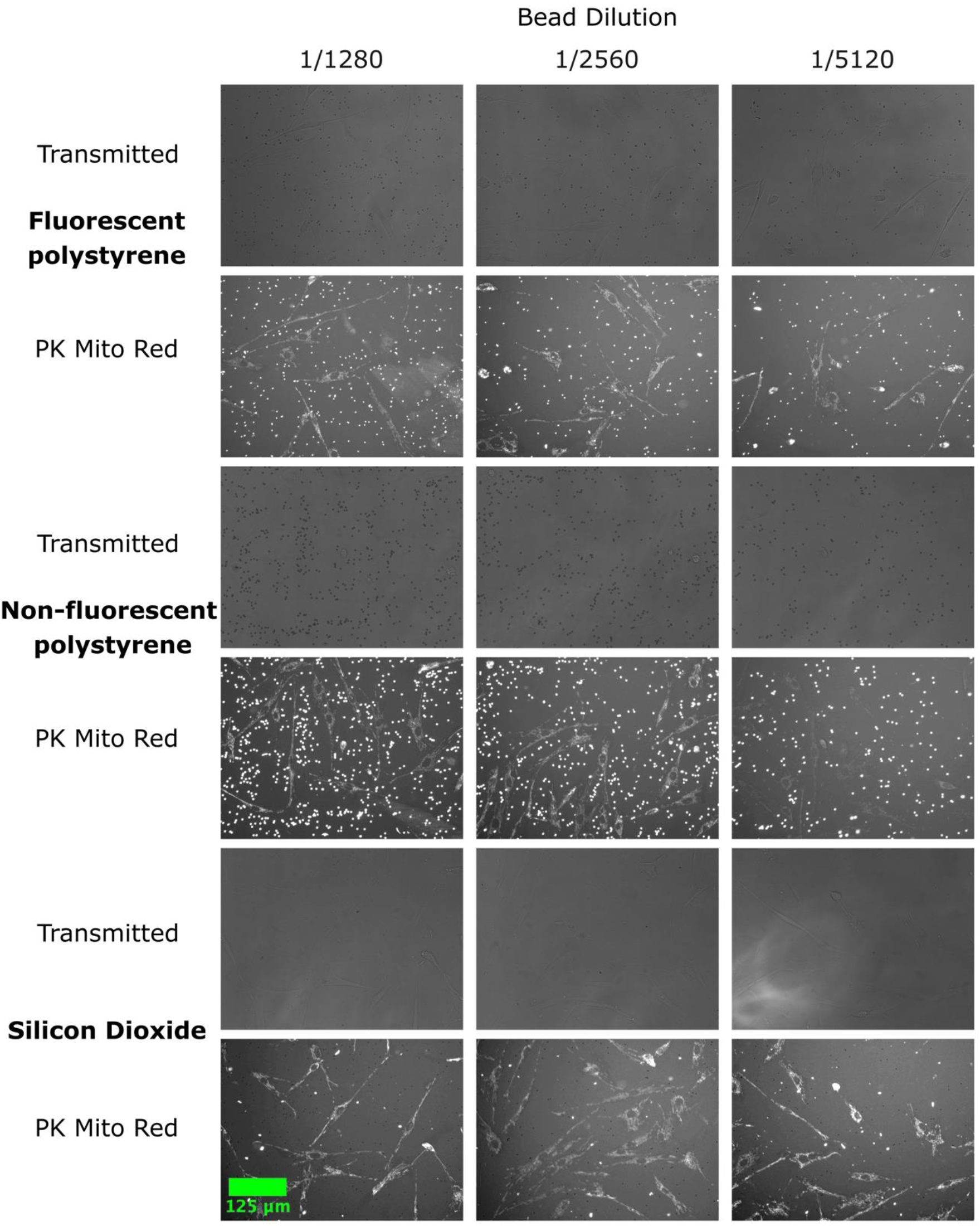
Example images of 3 distinct types of microbeads. Images have been acquired in experiment 1. Fluorescent (Texas Red) polystyrene (PS) 2μm, non-fluorescent polystyrene (PS) 3μm, and non-fluorescent silica dioxide 2μm microbeads were diluted to 1/1408, 1/2816 and 1/5632 as listed on the left) into wells of a 384 well plate containing fibroblasts stained with PK Mito far red and media. The plate was centrifuged at 1000 x g, for 5 minutes before imaging. Images were acquired using a 20X Plan fluorite objective on an EVOS M7000 microscope in brightfield and fluorescent mode. Autofocus was used in each channel. Scale bar = 125 um.

**Figure 7:**
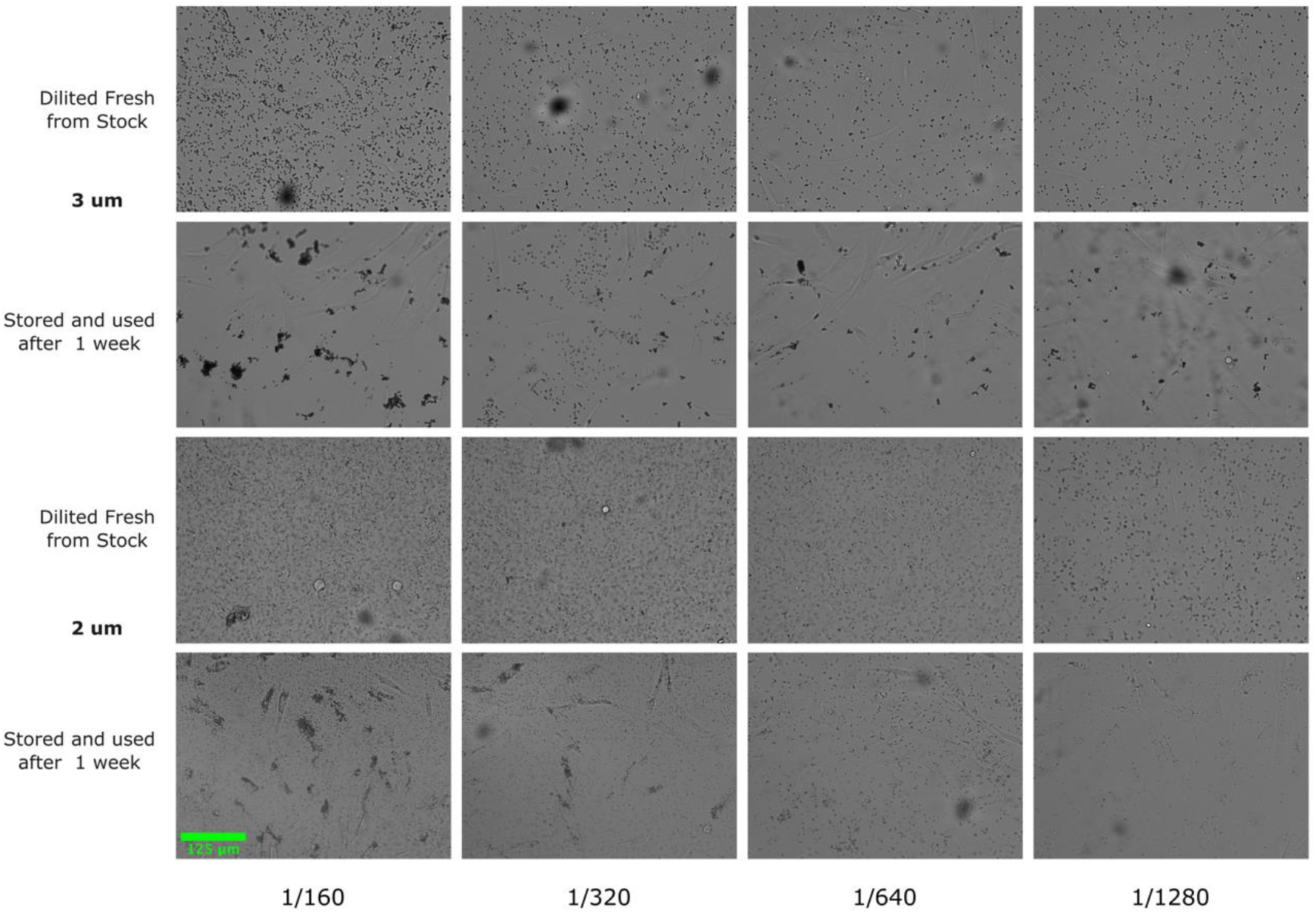
Beads diluted and stored in PBS aggregate over time. Images acquired on EVOS M7000 microscope using 20X Plan fluorite objective. Fluorescent (2 um) and non-fluorescent (3 um) beads were stored in 1.5 microcentrifuge tubes covered with foil at room temperature following dilution and are shown one week after storage. Results indicate beads should be diluted fresh with each experiment. Scale bar = 125 um.

## General notes and troubleshooting

### General notes

1. This protocol has been optimized for use with adherent cells.
2. In each experiment, the beads were diluted fresh. We found that diluting the beads and storing them in PBS resulted in aggregation (Figure below)
3. The exact procedure described above was used with the other beads evaluated in experiment 1. Beads were further diluted for experiment 1 than described in the procedure
4. In all experiments, a well with beads was imaged before wells without beads (Figure 2). If the design of experiment 2 will not be followed, it is good practice to decide where the wells with beads will go beforehand, to ensure this can be done based on scanning options available on the microscope.
5. Autofocus was implemented by the EVOS software which is used to control the microscope. The autofocus protocol described in the data acquisition section of the protocol was created using the Automate feature available on the EVOS software. The autofocus algorithm selected for all acquisitions is called fluorescence optimized. The algorithm is described in the EVOS manual: “the focal plane is derived from the highest ratio between detailed, high-contrast objects against the background”
6. This method has been evaluated solely on the EVOS imaging system.

